# The MuSK-BMP pathway maintains myofiber size in slow muscle through regulation of Akt-mTOR signaling

**DOI:** 10.1101/2022.11.05.514105

**Authors:** Diego Jaime, Lauren A. Fish, Laura A. Madigan, Madison D. Ewing, Justin R. Fallon

## Abstract

Myofiber size regulation is critical in health, disease, and aging. MuSK (muscle-specific kinase) is a BMP (bone morphogenetic protein) co-receptor that promotes and shapes BMP signaling. MuSK is expressed at all neuromuscular junctions and is also present extrasynaptically in the slow soleus muscle. To investigate the role of the MuSK-BMP pathway in vivo we generated mice lacking the BMP-binding MuSK Ig3 domain. These ΔIg3-MuSK mice are viable and fertile with innervation levels comparable to wild type. In 3-month-old mice myofibers are smaller in the slow soleus, but not in the fast tibialis anterior (TA). Transcriptomic analysis revealed soleus-selective decreases in RNA metabolism and protein synthesis pathways as well as dysregulation of IGF1-Akt-mTOR pathway components. Biochemical analysis showed that Akt-mTOR signaling is reduced in soleus but not TA. We propose that the MuSK-BMP pathway acts extrasynaptically to maintain myofiber size in slow muscle by promoting protein synthetic pathways including IGF1-Akt-mTOR signaling. These results reveal a novel mechanism for regulating myofiber size in slow muscle and introduce the MuSK-BMP pathway as a target for promoting muscle growth and combatting atrophy.

## INTRODUCTION

Maintaining myofiber size is essential for proper muscle function. Muscle atrophy characterizes aging, disuse, cancer cachexia and disease^1–3^. Notably, individual muscles are differentially affected in many of these settings^4^. For example, in sarcopenia upper leg muscles atrophy while the soleus muscle in the lower leg is spared^5^. In Duchenne muscular dystrophy limb-girdle muscles are affected in the first years of life while upper limb muscles are spared until several years later ^6^. In contrast, muscle weakness preferentially affects muscles in the anterior compartments of the face and leg in FSHD^4^. However, the molecular basis for such muscle-selective vulnerability to atrophy is largely unknown.

Members of the TGF-β (transforming growth factor-beta) superfamily, including myostatin and BMPs, are potent regulators of muscle size. Myostatin is a negative regulator of muscle mass, and its genetic deletion or pharmacological inhibition results in muscle hypertrophy^7^. In contrast, BMP signaling promotes muscle growth. Overexpression of BMP7 or a constitutively active BMP receptor BMPR1a (ALK3) in skeletal muscle results in increased muscle mass, fiber size, and elevated canonical BMP and Akt/mTOR signaling^8^. Inhibiting BMP signaling by overexpressing the BMP sequestering protein noggin abolishes the hypertrophic phenotype observed in myostatin-deficient mice^9^. Either increasing BMP or reducing myostatin signaling can restore myofiber size in cancer cachexia^2,10^. These studies suggest that the BMP and myostatin/activin pathways antagonize each other and tipping the balance can result in either hypertrophy or atrophy. They also implicate BMP signaling as an attractive target for combatting atrophy. However, targeting BMPs therapeutically is challenging since unlike myostatin, BMPs are expressed ubiquitously and serve a wide range of functions throughout the body^11,12^.

Recently, we discovered that MuSK (muscle-specific kinase) is a BMP co-receptor that promotes and shapes the BMP-induced transcriptional output in myogenic cells^13^. MuSK binds BMPs 2, 4 and 7 with low nanomolar affinity as well as the type I BMP receptors BMPR1a and 1b. Importantly, the MuSK Ig3 domain is necessary for high affinity BMP binding. Studies in cell culture show that MuSK enhances BMP4-induced pSmad1/5/8 signaling and that the expression of a large subset of BMP4-responsive transcripts is MuSK-dependent. Notably, this regulation requires the activity of Type I BMP receptors, but not that of the MuSK tyrosine kinase. Thus, the function of MuSK as a BMP co-receptor, which we term the MuSK-BMP pathway, is structurally and functionally distinct from its role in agrin-LRP4 signaling, which is essential for synapse formation, is mediated by the MuSK Ig1 domain, and absolutely requires MuSK tyrosine kinase activity^14,15^.

The function of the MuSK-BMP pathway in regulating myofiber size in vivo is unknown. However, several observations suggest that this pathway may be particularly important in slow (e.g., soleus) as compared to fast muscles (e.g. EDL and TA). For example, while MuSK is localized at all NMJs, it is also localized extrasynaptically in soleus, but not TA^16^. Further, single nuclei sequencing has shown that MuSK transcripts are expressed at elevated levels in soleus as compared to TA myonuclei, which is consistent with the 3-5 fold higher levels of MuSK in soleus compared to fast muscle^13,17^.

Here we investigated the role of the MuSK-BMP pathway in fast and slow muscle. To selectively manipulate the MuSK-BMP pathway, we took advantage of previous work showing that the MuSK Ig3 domain is necessary for high affinity BMP binding^13^, but dispensable for agrin-LRP4 binding and AChR clustering^18–20^. We generated mice where the MuSK Ig3 domain was deleted (ΔIg3-MuSK). These mice are fertile and viable with normal innervation levels. Cultured ΔIg3-MuSK myoblasts show reduced BMP signaling as judged by BMP-induced pSmad1/5 and gene expression. In vivo, ΔIg3-MuSK soleus muscle fibers were atrophied while TA fibers were unaffected. RNA-seq revealed largely non-overlapping changes in transcriptomic profiles and biological GO pathways and dysregulation of IGF1-Akt-mTOR components in ΔIg3-MuSK soleus compared to TA. Notably, in soleus multiple GO pathways involving RNA metabolism were selectively downregulated. The soleus also showed extracellular matrix (ECM) remodeling. Biochemical analysis revealed reduction of p4EBP1, a key component of the mTOR pathway, in soleus but not TA. We propose that the MuSK-BMP pathway promotes protein synthesis and maintains myofiber size in slow muscle via the Akt-mTOR pathway.

## RESULTS

### Generation of ΔIg3-MuSK mice

The MuSK Ig3 domain is necessary for high affinity BMP4 binding but is dispensable for agrin-LRP4 binding and AChR clustering^13,14,19^. To selectively perturb the MuSK-BMP pathway we generated a mouse model expressing an allele lacking the MuSK Ig3 domain (ΔIg3-MuSK). We used CRISPR-Cas9 to delete a ∼11 kb genomic region containing exons 6 and 7, which encode the MuSK Ig3 domain, to create the MuSK^ΔIg3^ allele (Fig. 1A, B). PCR amplification of the intronic regions of exon 5-6 and 5-8 borders yielded amplicons of the predicted size for the WT MuSK and MuSK^ΔIg3^ alleles, respectively (Fig. 1C). MuSK^ΔIg3/ΔIg3^ mice are viable and fertile with normal weights (Fig. 1D), grip strength (Fig. 1E), and innervation levels in both the fast sternomastoid (Fg. 1F) and the slow soleus (Fig. 1G). In this study we will term the MuSK^ΔIg3/ΔIg3^ animals ‘ΔIg3-MuSK’.

**Figure 1.**
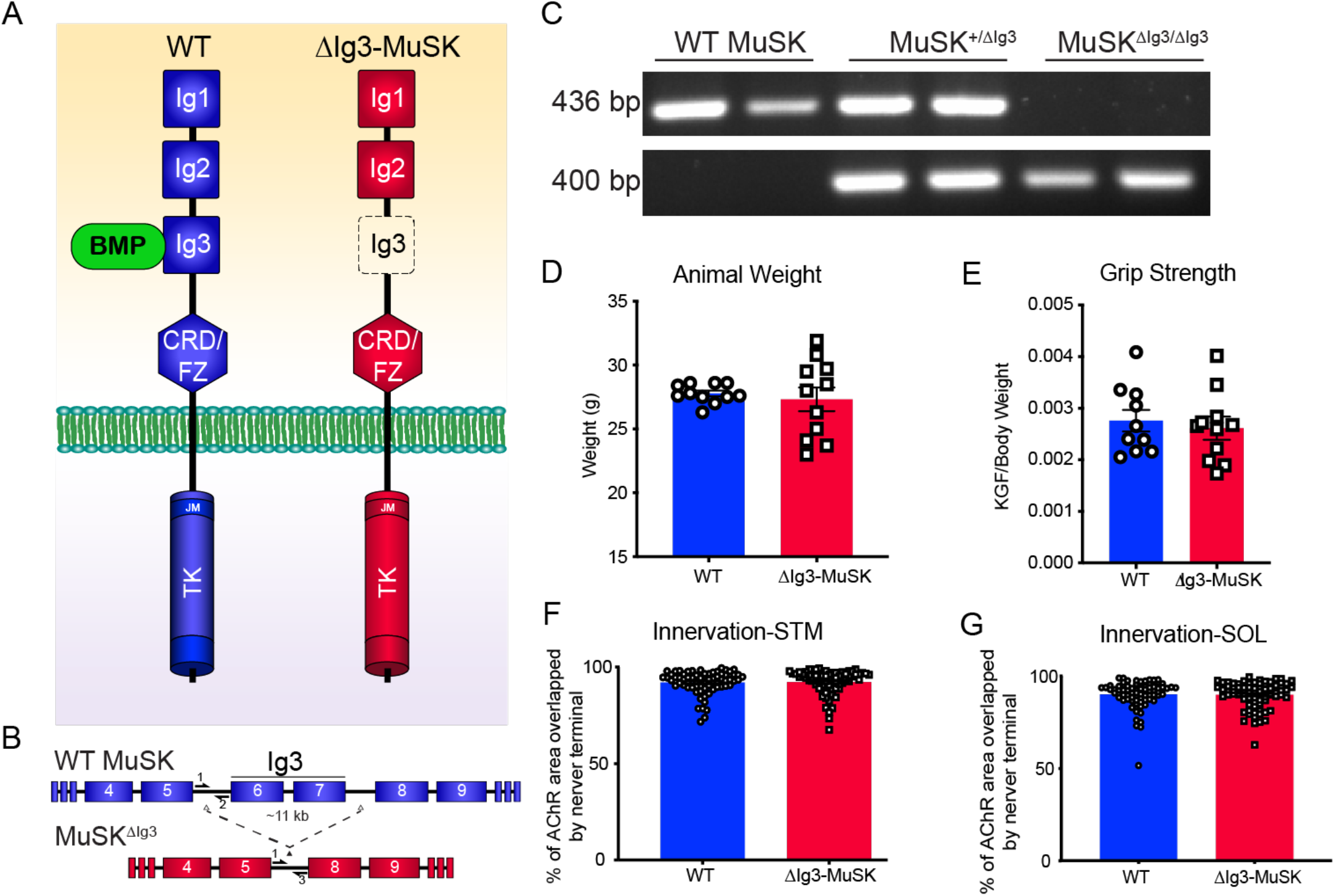
Generation of the ΔIg3-MuSK mouse. (A) Schematic representation of full length and ΔIg3-MuSK isoforms. The extracellular full-length isoform contains three immunoglobulin (Ig)-like domains (Ig1, Ig2, and Ig3) as well as a CRD/Fz and an intracellular tyrosine kinase domain. ΔIg3-MuSK lacks the BMP-binding Ig3 domain. (B) CRISPR-Cas9 was used to delete exon 6 and 7, which encode the Ig3 domain, along with the intervening intronic sequence to generate the ΔIg3-MuSK allele. (C) PCR amplification of WT and ΔIg3-MuSK alleles in WT and ΔIg3-MuSK heterozygous and homozygous mice using allele-specific primers (depicted in B). The animal weights (D) and grip strength (E) of 3-month-old WT and ΔIg3-MuSK were comparable. Innervation in the Ig3-MuSK animals, as assessed by overlap of pre- and post-synaptic structures, was equivalent to that in WT in both the fast sternomastoid (F) and slow soleus (G) (n=6 animals per genotype).

### BMP4 signaling is perturbed in ΔIg3-MuSK cells

We first generated stable ΔIg3-MuSK myogenic cell lines to probe MuSK expression and BMP signaling in cells lacking the MuSK Ig3 domain (see Methods). Immunostaining of unpermeabilized myoblasts with a monoclonal antibody directed against the MuSK Ig2 domain showed that MuSK is expressed at comparable levels and distribution at the cell surface in both WT and ΔIg3-MuSK myoblasts (Fig. 2A). MuSK transcript levels are also comparable in WT and ΔIg3-MuSK cells (Fig. 2B). To probe the role of the MuSK Ig3 domain in the BMP4 response, we serum-starved WT and ΔIg3-MuSK myoblasts and examined the nuclear localization of pSmad1/5 in the absence or presence of added BMP4. As expected, nuclear pSmad1/5 levels were low at baseline in both genotypes. However, following BMP4 stimulation the level of nuclear pSmad1/5 was higher in WT compared to ΔIg3-MuSK cells (Fig. 2C). We then performed Western blots to quantify the pSmad1/5 response to BMP4 treatment (Fig. 2D). The ΔIg3-MuSK cells showed a reduction in BMP4-stimulated pSmad1/5 levels compared to WT (Fig. 2D, E).

**Figure 2.**
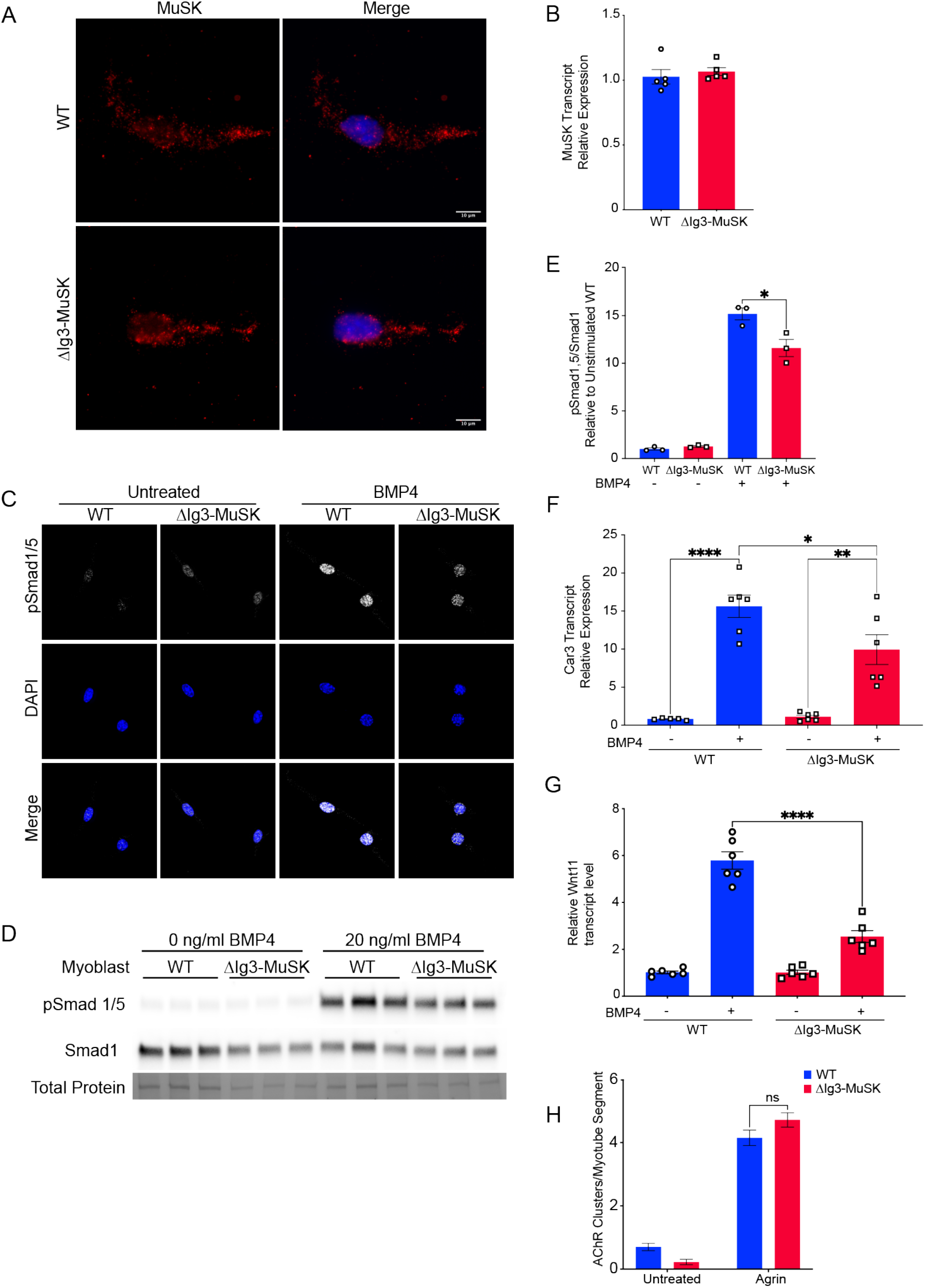
BMP signaling is selectively perturbed in ΔIg3-MuSK myogenic cells. (A, B), MuSK expression: (A) Unpermeabilized WT and ΔIg3-MuSK myoblasts cells were fixed, and cell surface MuSK was visualized by immunostaining with an antibody directed against the MuSK Ig2 domain (red). Note that comparable levels and distribution of immunostaining were observed in both genotypes. (B) WT and ΔIg3-MuSK myoblast cell lines express comparable levels of MuSK transcript as assessed by qRT-PCR. Data are means ± SEM from five biological replicates and two independent experiments. (C-G), BMP signaling: (C) WT and ΔIg3-MuSK myoblasts were treated with 20 ng/ml BMP4 for 15 min and immunostained for pSmad1/5. Note the increased intensity of pSmad staining in nuclei of BMP4-treated WT compared to ΔIg3-MuSK cells. Data are from three independent experiments with three biological replicates per group. (D) pSmad1/5 and total Smad1 in BMP4-treated cells (as in C) were assessed by Western blotting. (E) The level of BMP4-induced pSmad1/5, normalized to total protein, was reduced ΔIg3-MuSK as compared to WT cells. Data are means ± SEM of three biological replicates and independent Western blots (**** p < 0.0001, two-way ANOVA with Bonferroni’s multiple comparisons). (F) Cells were treated as in (C) and Car3 transcript levels were assessed. Data are mean ± SEM of 5-6 biological replicates per condition and replicated twice (**** p < 0.0001, *** p < 0.001, * p < 0.05, two-way ANOVA with Bonferroni’s multiple comparisons). (G) Cultured primary myotubes from WT and ΔIg3-MuSK mice were treated with 25 ng/ml BMP4 for 2 hr and levels of Wnt11 mRNA were measured by qPCR. Note that ΔIg3-MuSK myotubes show reduced Wnt11 expression in response to BMP4 stimulation. Data are means ± SEM from 6 biological replicates from two experiments. (**** p < 0.0001, two-way ANOVA with Bonferroni’s multiple comparisons). (H) Agrin signaling: WT and ΔIg3-MuSK cultured primary myotubes were treated with agrin for 16 hours. Acetylcholine receptors (AChR) were stained with α-bungarotoxin (red) and number of AChR clusters per myotube were counted (two-way ANOVA with Bonferroni’s multiple comparisons).

Since BMP4 treatment induces the expression of the MuSK-dependent genes Car3 and Wnt11^13^, we then assessed the role of the MuSK Ig3 domain in regulating these BMP4-induced transcripts. Fig. 2F shows that levels of BMP4-induced Car3 transcripts are lower in ΔIg3-MuSK myoblasts compared to WT. Similarly, BMP signaling is also perturbed in primary cultured myotubes, where BMP4-induced Wnt11 transcript levels were lower in ΔIg3-MuSK compared to WT myotubes (Fig. 2G). Finally, to establish that agrin signaling is preserved in ΔIg3-MuSK cells, we assessed agrin-induced AChR clustering in WT and ΔIg3-MuSK primary myotubes and found that the level of agrin-induced AChR clustering was comparable in both genotypes (Fig. 2H).

This normal agrin response in cultured cells is in agreement with the comparable innervation levels observed in WT and ΔIg3-MuSK muscle (Fig. 1G, H). Taken together, these data show that in cultured cells the MuSK Ig3 domain regulates BMP4 pSmad1/5 and transcriptional responses and that agrin- and MuSK-BMP-dependent signaling can be clearly distinguished.

### Unique transcriptional profiles in ΔIg3-MuSK fast and slow muscles

The results from cultured ΔIg3-MuSK cells indicated that the MuSK Ig3 domain regulates BMP4 signaling and transcriptional response. We next explored the role of the MuSK-BMP pathway in vivo. The soleus muscle expresses several fold higher levels of MuSK transcript compared to the TA^13,16^, suggesting that MuSK may play a particularly important role in the slow soleus. We compared the transcriptional profiles in 3-month-old WT and ΔIg3-MuSK TA and soleus using RNA-seq. We isolated RNA from 36 muscles (n=9 TA and soleus muscles per genotype) and sequenced at a depth of ∼50 million reads/sample. The results revealed striking differences in the transcriptional profiles between these muscles. Figs. 3A and 3B show heatmaps of the transcriptomic profile differences in the 1000 most variable genes in ΔIg3-MuSK compared to WT in TA and soleus, respectively. Principal component analysis of gene expression profiles showed that both the ΔIg3-MuSK TA and soleus muscles clustered separately from WT (Fig. 3C, D). Analysis of differentially expressed genes (DEGs) revealed significant changes in both muscles (Table S1, S2). The TA showed 485 upregulated and 287 downregulated genes compared to WT (Fig. 3E), while the soleus exhibited 531 upregulated and 253 downregulated genes (Fig. 3F). Notably, although the absolute number of DEGs was similar in both muscles, the changes in soleus showed broader ranges of log2 fold change and log10 FDR values compared to TA (Fig. 3G, H), indicating that the magnitude and significance of the DEG changes were larger in the soleus compared to the TA. Finally, the identity of DEGs in TA and soleus were strikingly different: only 19/799 of the combined downregulated genes and 34/704 of the combined upregulated genes were shared between TA and soleus (Fig. 3l). Taken together, these results show that the transcriptional profiles of TA and soleus WT and ΔIg3-MuSK muscles are qualitatively and quantitatively distinct, with the soleus transcriptome being more affected by the downregulation of MuSK-BMP signaling.

**Figure 3.**
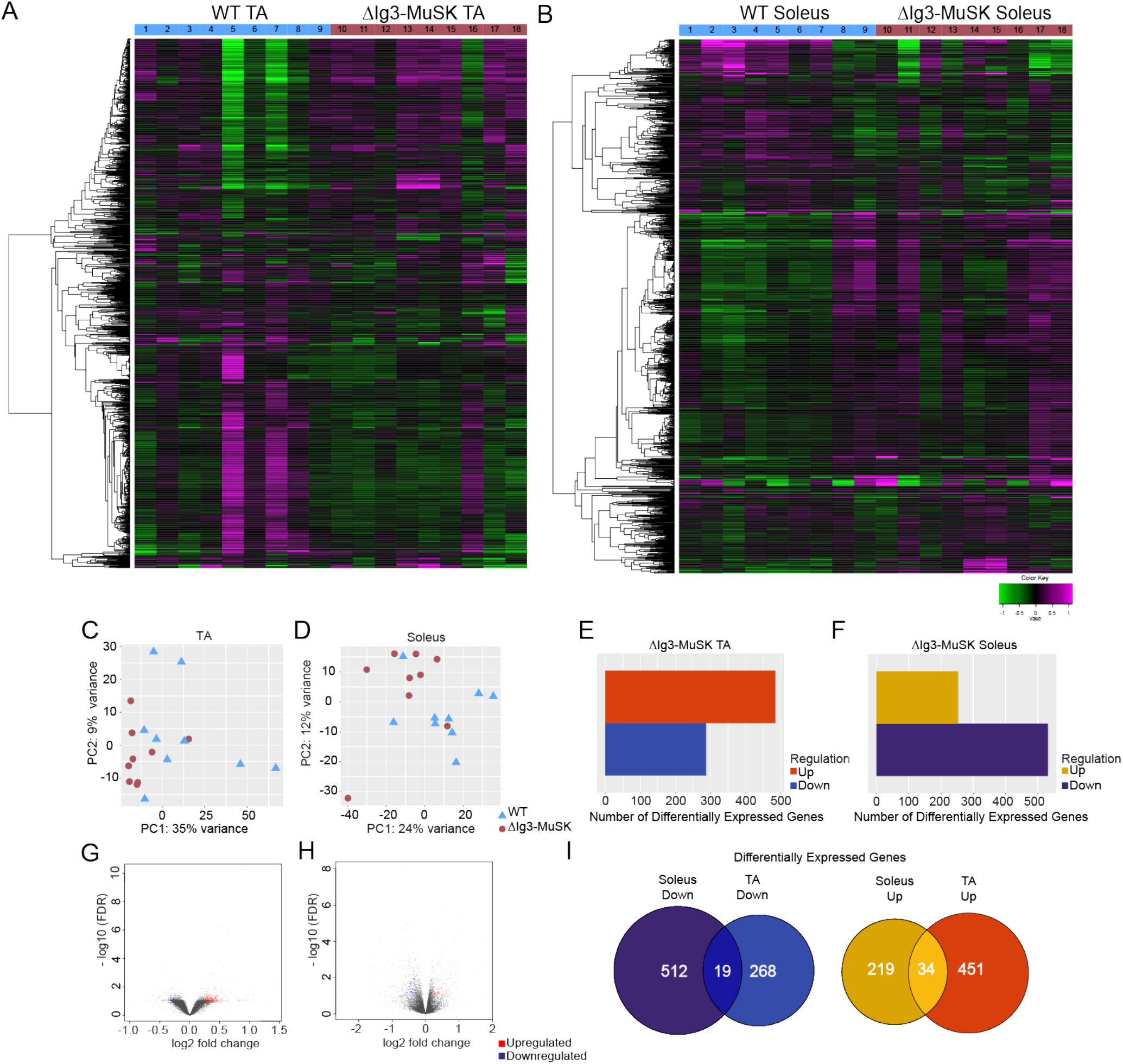
Distinct transcriptomic profiles in ΔIg3-MuSK TA and soleus. WT and ΔIg3-MuSK soleus and TA muscles were subjected to RNA-sequencing at a depth of ∼50 million reads per sample (n = 9 muscles/genotype). Heat map of the top 1000 variable genes in WT and ΔIg3-MuSK TA (A) and soleus (B). Each column represents one muscle sample. Principal component analysis of WT and ΔIg3-MuSK TA (C) and soleus (D) shows clustering of samples with similar gene expression profiles (▲, WT; ●, ΔIg3-MuSK; individual muscles). (E, F) Differentially expressed genes (DEGs) with 1.2 minimum fold-change and 0.1 false discovery rate (FDR) were determined using DEseq2 (See also Table. S1, S2). (G, H) Volcano plot of -log10 FDR versus gene log2 fold change (red dots up DEGs; blue dots down DEGs). (I) Venn diagram depicting the shared and unique upregulated and downregulated DEGs between ΔIg3-MuSK TA and soleus.

To gain insight into cellular mechanisms impacted in ΔIg3-MuSK muscles, we performed gene set enrichment analysis (GSEA) using GO terms for both TA and soleus. The specific pathways and the extent of change within pathways differed markedly in TA and SOL (Fig. 4A and B). Multiple upregulated pathways in the TA were related to regulation of GTPase activity and Ras/Rho signal transduction (Fig. 4A, Table S3). In contrast, in the soleus the major upregulated pathways were related to ECM structure, ECM organization, and inflammation, suggesting muscle damage (Fig. 4B, Table S3). In addition, we found upregulated pathways related to ERK1/2 signaling regulation as well as collagen fibril organization (Fig. 4B). Pathways downregulated in the TA included those related to translation, peptide biosynthesis, protein folding and ribosome biogenesis (Fig. 4A). Additionally, there were multiple processes related to mitochondrial energy metabolism such as mitochondrial organization, respiratory transport chain, mitochondrial ATP synthesis, and proton transport (Fig. 4A). The soleus showed a striking downregulation of biological process pathways related to RNA metabolism including mRNA and ncRNA processing, RNA splicing, and ribonucleoprotein complex biogenesis and subunit organization (all adjusted p-values between 10^−6^ and 10^−11^). These changes suggest that the ΔIg3-MuSK soleus exhibits dysfunctional post-transcriptional RNA metabolism and protein synthesis (Fig. 4B). Finally, the GO pathways also showed striking statistical differences in TA vs soleus. In soleus a cluster of down RNA-related pathways had p values between 10^−7^ and 10^−12^, while all the TA values (up or down) were ≥10^−7^.

**Figure 4.**
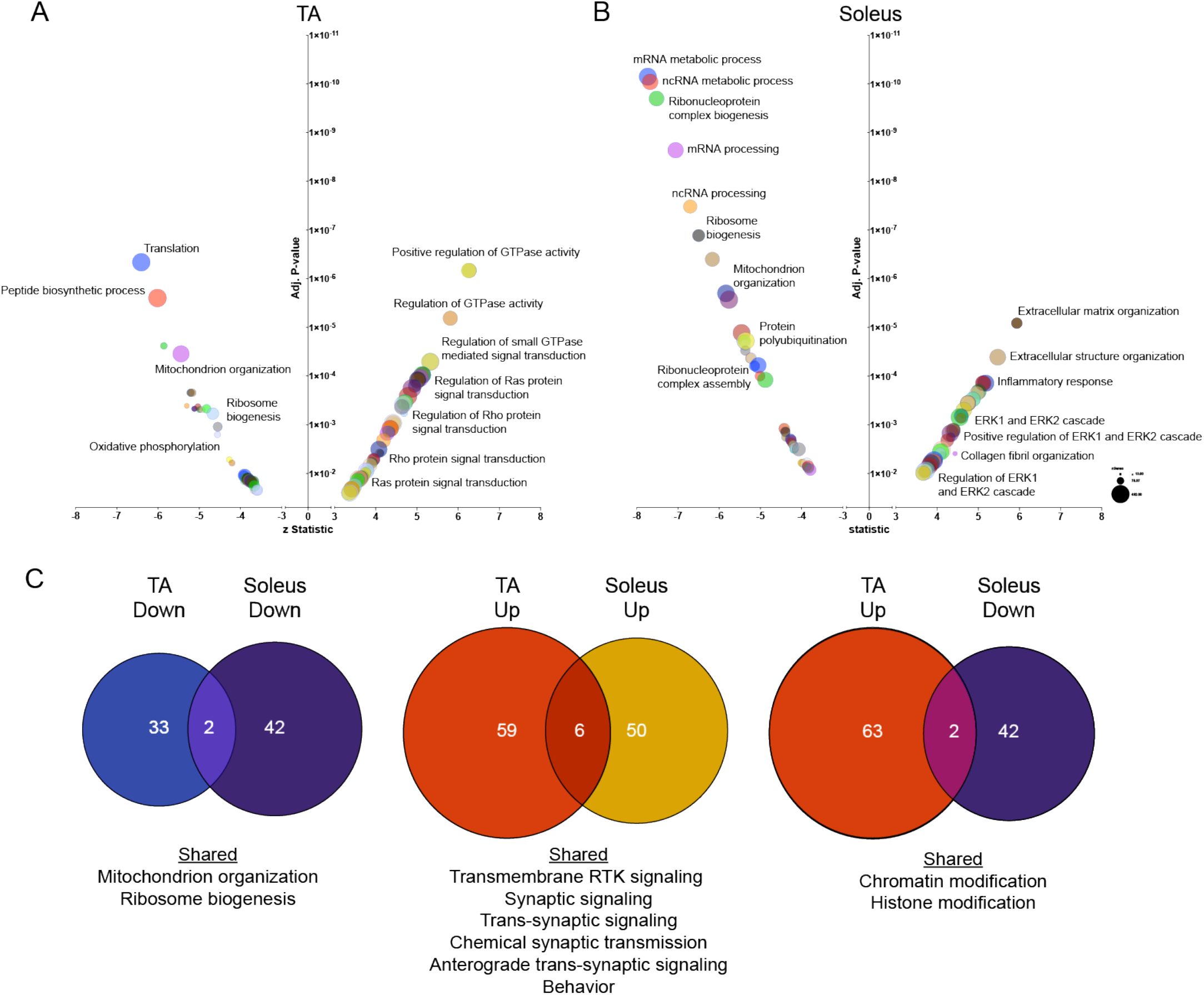
Distinct pathways are dysregulated in ΔIg3-MuSK TA and soleus. (A, B) Gene ontology pathway analysis of top 100 upregulated and downregulated biological process pathways in ΔIg3-MuSK TA (A) and soleus (B) (0.1 FDR; See Table S3). Note the number of highly significant pathways in soleus (adjusted p values between 10^−7^ and 10^−12^) compared to TA. Selective enrichment in soleus down pathways were related to translation including those involving ribosome biogenesis as well as mRNA and ncRNA metabolism. Soleus up pathways included those involved in ECM and inflammation. (C) Venn diagram showing the number of shared and unique downregulated and upregulated pathways between ΔIg3-MuSK TA and soleus (see Table S3). Note that ≤5% of pathways are shared in any of the comparisons. No shared pathways were observed when comparing TA down and soleus up.

Our analysis also revealed a small number (<5%) of shared GO pathways in TA and soleus, including downregulation of mitochondrion organization and ribosome biogenesis (Fig. 4C). Shared upregulated pathways included synaptic signaling, trans-synaptic signaling, chemical synaptic transmission and anterograde trans-synaptic signaling, suggesting that the MuSK-BMP pathway may also play a role at the neuromuscular junction (Fig. 4C). Some pathways showed opposite directionality, with chromatin organization and histone modification downregulated in soleus while upregulated in the TA (Fig. 4C). These analyses suggest that the MuSK-Ig3 domain regulates distinct pathways in ΔIg3-MuSK TA and soleus.

### Myofiber size is reduced in ΔIg3-MuSK soleus

The muscle-selective reductions in RNA metabolism pathways and ribosome biogenesis raised the possibility that myofiber size is reduced in ΔIg3-MuSK soleus. The overall structure of both the TA and the soleus as revealed by H&E staining was similar in both genotypes (Fig. 5A, B), with no evidence of degeneration observed. We did detect transcriptomic structural signatures related to increased ECM and inflammatory pathways (Fig. 4). While WGA staining in the TA was comparable in both genotypes (Fig. 5C), we observed a marked increase in WGA signal in ΔIg3-MuSK soleus (Fig. 5D). The staining levels for dystrophin were comparable in both muscles across genotypes (Fig. 5E, F). Muscle fiber Feret diameter analysis in TA revealed that the myofiber sizes were comparable in ΔIg3-MuSK and WT (Fig. 5G). In contrast, myofiber sizes were reduced in the ΔIg3-MuSK soleus (Fig. 5H). This myofiber atrophy was observed when all myofibers were scored (Fig. 5H) and when either type I (Fig. 5I) or type IIa (Fig. 5J) were separately analyzed using double labeling with myosin-specific markers (see Methods). Type I and Type IIa myofibers comprise approximately 80% of the myofibers in this muscle ^21^. Thus, the large majority of myofiber types in the soleus are atrophied, while the TA myofibers are unaffected in ΔIg3-MuSK mice at this age.

**Figure 5.**
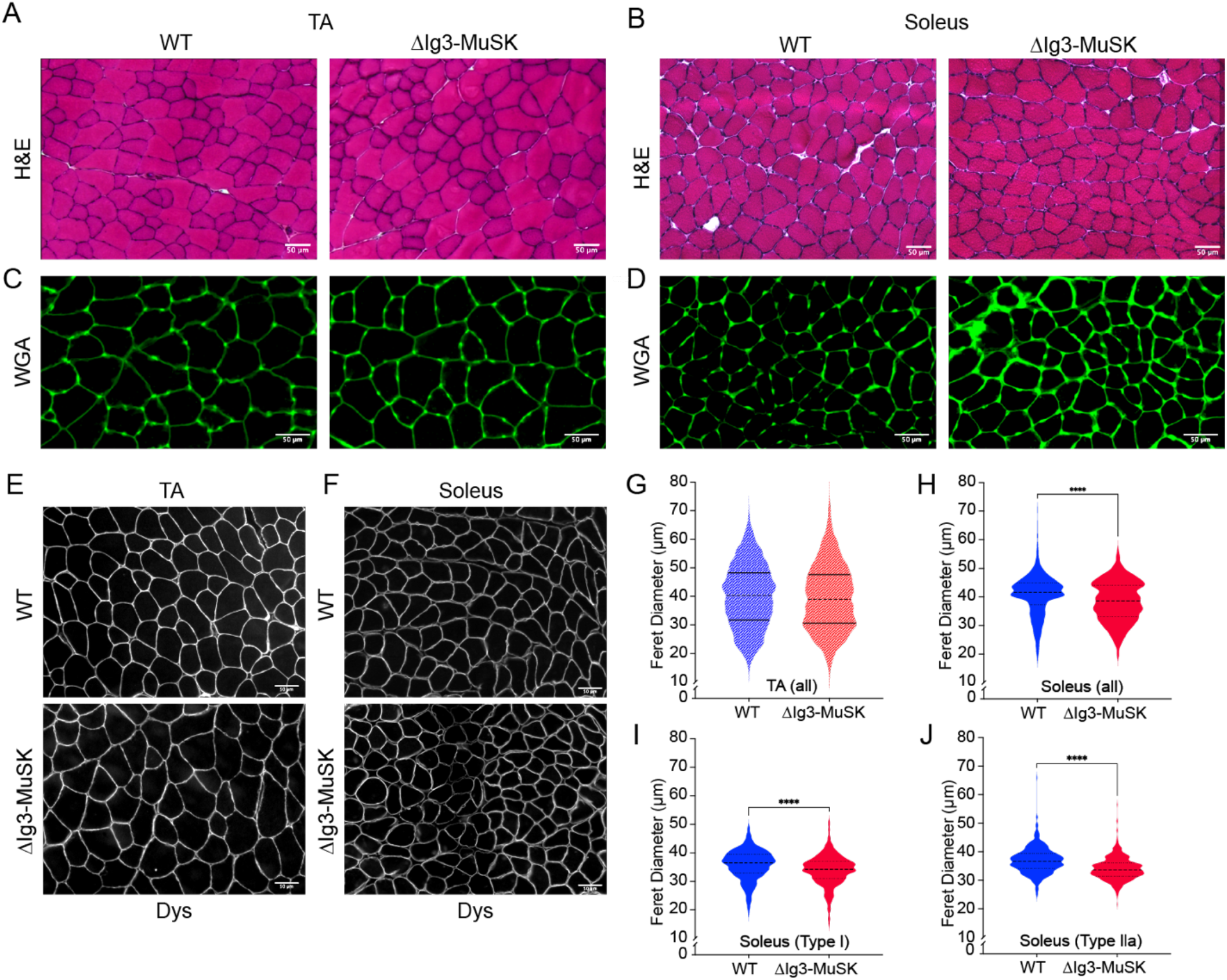
Reduced myofiber size in ΔIg3-MuSK soleus but not TA. WT and ΔIg3-MuSK TA and soleus sections were stained with H&E (A, B), WGA (C, D), or anti-dystrophin (E, F). In the TA, the structure (A), WGA staining (C), and mean Feret diameters (G), were indistinguishable in both genotypes. Data are means ± SEM from three animals of each genotype. Average number of muscle fibers counted per animal 963 ± 77. Soleus WT and ΔIg3-MuSK stained with H&E (B), WGA (D), or dystrophin (F). Note that the overall structure of the muscle was comparable, but there is increased interstitial WGA signal in ΔIg3-MuSK soleus. Soleus myofiber Feret diameters of all fibers (H), Type I fibers (I), or Type IIa fibers (J), were reduced in the ΔIg3-MuSK compared to WT. All fibers: 40.6 μm ± 0.2 WT and 38.4 μm ± 0.2 ΔIg3-MuSK (697 fibers/animal ± 72, n=3 animals); Type I fibers: 36.6 μm ± 0.2 WT and 34.4 μm ± 0.2 ΔIg3-MuSK (300 ± 33 fibers/animal, n=4 animals); Type IIa: 36.7 μm ± 0.2 WT and 33.9 μm ± 0.2 ΔIg3-MuSK (487 ± 50, fibers/animal n=4 animals; **** P < 0.0001, Mann-Whitney test).

### MuSK expression in WT and ΔIg3-MuSK muscle

We next examined MuSK mRNA expression and protein localization in WT and ΔIg3-MuSK muscles. We first assessed the levels of MuSK transcripts, which in cultured muscle cells is regulated by the MuSK-BMP pathway^13^. We and others have previously reported that MuSK transcripts are expressed at four to five-fold higher levels in the soleus compared to fast muscles such as EDL and TA^13,16^. We confirmed these findings in WT EDL and soleus (Fig. 6A). MuSK transcript levels were also higher in soleus as compared to EDL in ΔIg3-MuSK animals (Fig. 6B). MuSK expression in EDL was comparable in ΔIg3-MuSK and WT (Fig. 6C). However, MuSK transcript levels in soleus were reduced by ∼1/2 in ΔIg3-MuSK compared to WT (Fig. 6D).

**Figure 6.**
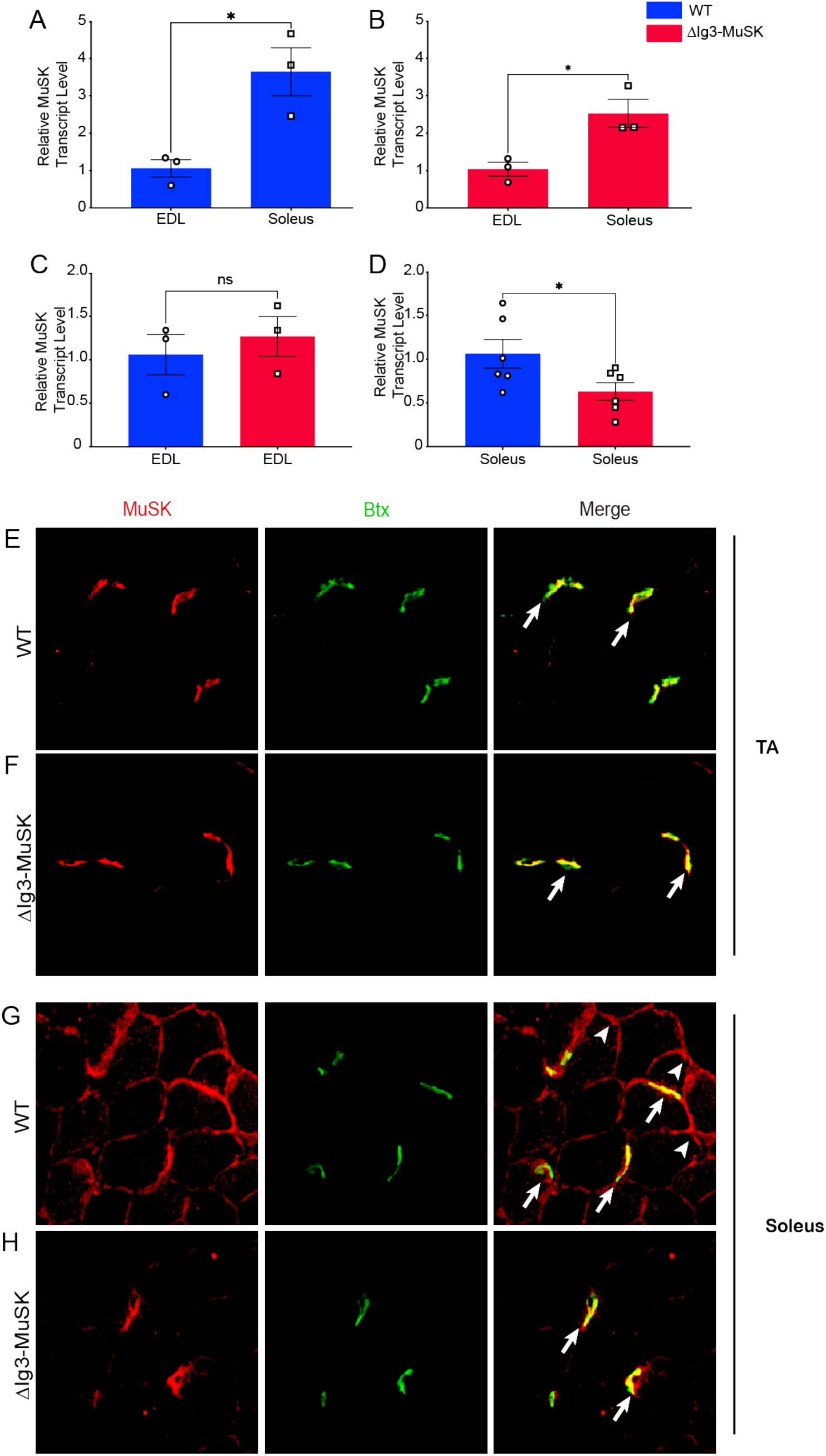
MuSK expression and extrasynaptic localization are reduced in ΔIg3-MuSK soleus. (A-D): MuSK transcript levels in WT and ΔIg3-MuSK EDL and soleus muscle were quantified by qRT-PCR. Note that MuSK transcript levels are selectively reduced in ΔIg3-MuSK soleus compared to WT while the levels in the EDL are comparable in both genotypes. Data are means ±SEM from 3 to 6 different biological replicates (*p < 0.05, unpaired two-tail Student’s t test). (E, F): MuSK localization in TA. TA sections from WT and ΔIg3-MuSK animals were immunolabeled with anti-MuSK (red), and α-bungarotoxin (green; to visualize NMJs). In the TA, MuSK is restricted to the NMJ (arrows) in both WT and ΔIg3-MuSK muscle. No extrajunctional MuSK localization is detected in TA of either genotype. (G, H): MuSK localization in soleus. In WT soleus (G) MuSK is localized at both the NMJ (arrows) and extrajunctionally (arrowheads). In contrast, in ΔIg3-MuSK soleus (H) MuSK is present at NMJs at levels comparable to WT, but extrajunctional MuSK was not observed.

We next assessed the localization of MuSK protein. MuSK was localized at all NMJs examined in both TA and soleus in WT and ΔIg3-MuSK muscle (Fig. 6E-H). In agreement with previous reports, in WT muscles MuSK localized at extrajunctional domains in the soleus but not in TA^16^. However, MuSK was not detected in extrajunctional domains in the ΔIg3-MuSK soleus (Fig. 6H). Taken together, these data indicate that MuSK-BMP signaling is important for regulating the expression and localization of extrajunctional MuSK in the soleus.

### Reduced Akt-mTOR signaling in ΔIg3-MuSK soleus

We next investigated potential signaling pathways that may mediate the selective atrophy of soleus fibers in ΔIg3-MuSK mice. The transcriptomic analysis (Fig. 4) showed that multiple RNA metabolism pathways were selectively downregulated in the ΔIg3-MuSK soleus. These observations suggested that reduced protein synthesis could underlie the atrophy observed in the soleus. One candidate pathway is the Akt-mTOR, which plays a central role in regulating muscle growth^22,23^. Moreover, several members of this pathway are selectively dysregulated in the ΔIg3-MuSK soleus including Igf1, Igf2bp2, Igfbp2, IRS1, Txnip and its antisense lncRNA Gm15441 (Supplemental Table 6). To directly assess the activity of the Akt-mTOR pathway in ΔIg3-MuSK mice, we measured the phosphorylation of 4EBP1, a major downstream target of mTOR (Fig. 7). Translation is promoted by phosphorylated 4EBP1 (p4EBP1) and is inhibited by the unphosphorylated form (4EBP1). We probed Western blots of muscle extracts from soleus and TA muscles of each genotype with antibodies against 4EBP1 and p4EBP1 (Fig. 7A and B,). Quantification showed that p4EBP1 levels were reduced by >40% in the ΔIg3-MuSK soleus (1 ± 0.15 and 0.55 ± 0.06, ± SEM in WT and ΔIg3-MuSK, respectively; p=0.02; n=5 muscles/genotype) (Fig. 7C). In contrast, p4EBP1 levels were comparable in WT and ΔIg3-MuSK TA (Fig. 7D). Notably, pSmad 1/5 levels did not differ in ΔIg3-MuSK compared to WT for both the TA and soleus (Fig. 7E and F). Taken together, these results indicate that the Akt-mTOR pathway is selectively reduced in ΔIg3-MuSK soleus

**Figure 7.**
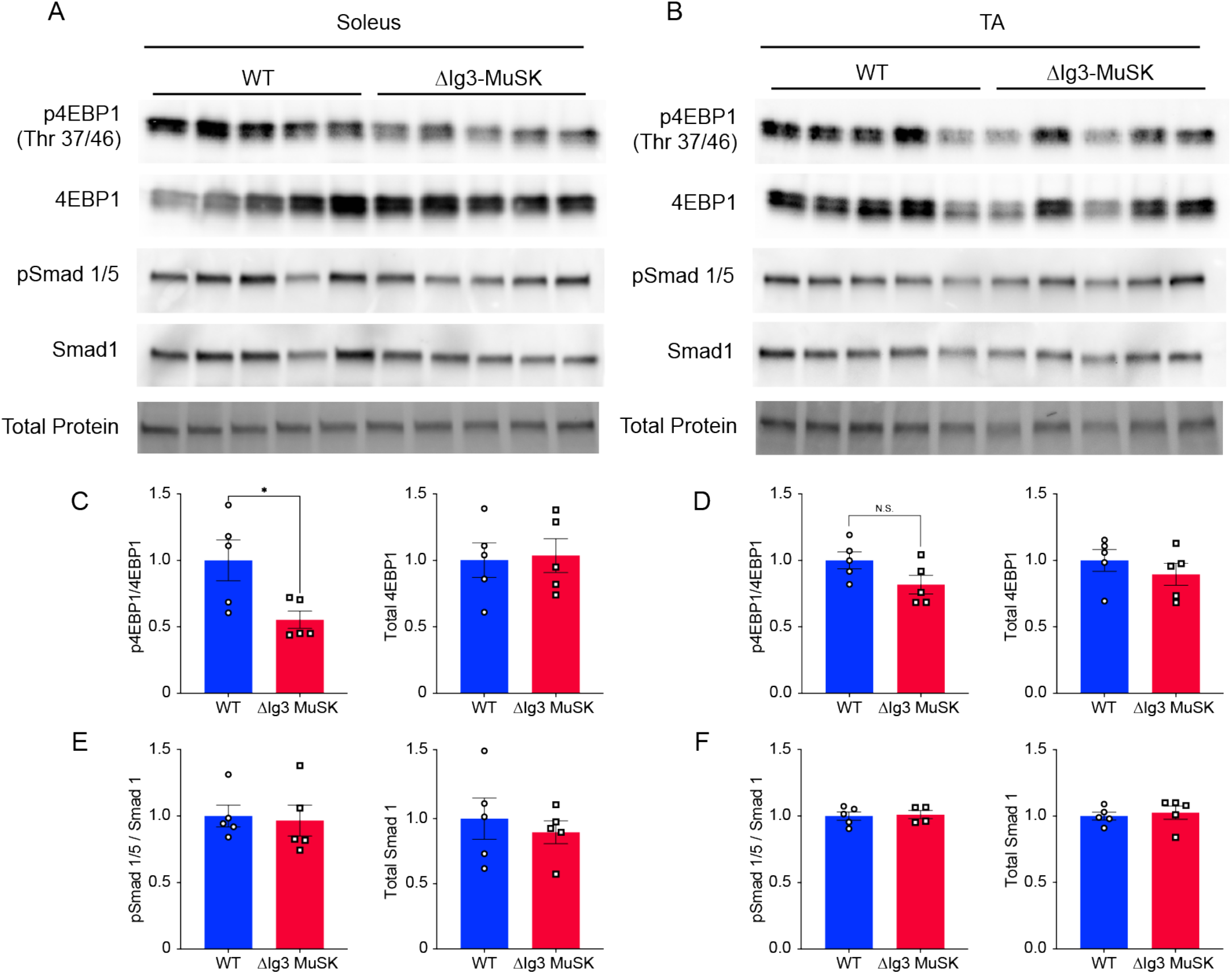
4EBP1 phosphorylation is reduced in ΔIg3-MuSK soleus but not TA. (A) WT and ΔIg3-MuSK soleus muscle protein extracts were isolated for Western blotting of pSmad1/5, total Smad1, p-4EBP1, and total 4EBP1. Total protein stain was used as loading control and for protein normalization. Levels of pSmad1/5 and p4EBP1 were determined as a ratio to total Smad1 and total 4EBP1, respectively. Note that p-4EBP1 levels were reduced in ΔIg3-MuSK soleus compared to WT (C), but unchanged in TA (D). pSmad 1/5 levels in ΔIg3-MuSK soleus and TA were comparable to WT (E and F). Data are means ± SEM from five biological replicates. Data are means ± SEM from five biological replicates (*p < 0.05, unpaired two tailed Student’s t test).

## DISCUSSION

In this report we introduce the MuSK-BMP pathway as a novel regulator of myofiber size in slow muscle in mice at 3-months of age. This pathway is selective for the slow soleus muscle as compared to the predominantly fast TA. In ΔIg3-MuSK mice, the soleus is atrophied, RNA metabolic pathways are downregulated and Akt-mTOR signaling is reduced compared to TA. Our findings indicate that the locus of MuSK-BMP action is extrasynaptic throughout the soleus, rather than secondary to changes at the NMJ. These results also reveal a novel in vivo function for MuSK that is independent of its role in agrin-LRP4 signaling at the synapse. The results shed light on the mechanism for the muscle-selective regulation of myofiber size and reveal a new pathway for promoting muscle growth and combatting atrophy.

Our results demonstrate that deletion of the MuSK Ig3 domain selectively perturbs MuSK function as a BMP co-receptor. First, MuSK protein lacking the Ig3 domain is localized at the cell surface in cultured ΔIg3-MuSK myogenic cells and the levels of MuSK mRNA are comparable in ΔIg3-MuSK and WT cells (Fig. 2). ΔIg3-MuSK is also localized at all NMJs examined in vivo (Fig 6; see further discussion below). This high-fidelity expression and localization are consistent with the fact that the ΔIg3-MuSK allele created by gene editing mimics a natural MuSK splice isoform^20,24^. Second, cultured ΔIg3-MuSK myogenic cells show reduced levels of pSmad 1/5 signaling and target gene expression in response to BMP treatment (Fig. 2). Notably, the MuSK-regulated BMP induced transcripts include Wnt11 and Car3, which were previously identified in a study using cultured MuSK^-/-^ myogenic cells^13^. In contrast, the agrin-LRP4 mediated functions of MuSK, which require its Ig1 domain, are spared: agrin-induced AChR clustering is robust in cultured ΔIg3-MuSK myotubes; in vivo, NMJs form normally, and innervation levels and grip strength are comparable to WT (Fig. 1).

Our transcriptomic, morphological, and biochemical results show that the MuSK-BMP pathway plays a selective role in slow as compared to fast muscle. The MuSK-BMP pathway regulates myofiber size in slow muscle. In soleus, muscle atrophy was observed in both type I as well as IIa fibers (Fig. 5), which together comprise ∼80% of fiber types in soleus. In contrast, no significant differences were observed in TA myofiber diameter in the 3-month age animals examined here (Fig. 5). The sets of both the up- and down-regulated genes in soleus and TA were also remarkably distinct. Out of a total of 1503 DEGs, only 19 downregulated and 34 upregulated genes were shared between soleus and TA, respectively (Fig. 3). GO pathway analysis also revealed distinct functions for the MuSK-BMP pathway in soleus and TA (Fig. 4; Table S3). As discussed below, a large number of downregulated GO pathways involved in RNA metabolism were unique to soleus. However, it is noteworthy that the two shared downregulated GO pathways were related to mitochondria organization and ribosome biogenesis, which raises the possibility that the MuSK-BMP pathway may regulate energy metabolism and some aspects of protein synthesis in both fast and slow muscle. Finally, we observed a number of pathways related to synaptic signaling and organization, raising the possibility that the MuSK-BMP pathway, while not essential for synapse formation, may play a role at the NMJ.

Several lines of evidence indicate that the MuSK-BMP pathway maintains soleus myofiber size through the regulation of the IGF1-Akt-mTOR pathway, the primary anabolic regulator of muscle cell size (Fig. 8)^22,25–30^. This pathway increases protein synthesis through mTOR-mediated phosphorylation of key elements regulating translation, notably 4EBP1. Our transcriptomic analysis revealed a host of downregulated GO pathways in RNA metabolism as well as dysregulation of members of the IGF1-Akt-mTOR pathway that were selective for the soleus. Importantly, biochemical analysis showed that p4EBP1, a direct target of mTOR, is downregulated in ΔIg3-MuSK soleus but not TA (Fig. 7). In agreement with our results, others have observed muscle-selective effects of rapamycin treatment on regulating myofiber size^29^. We saw no evidence that this atrophy was due to denervation, since the NMJs in both muscles were fully innervated (Fig. 1). Further, our transcriptomic analysis detected few signatures of upregulated protein degradation, such as atrogenes^31^ (Table S4, S5). Taken together, our results support a model where the MuSK-BMP pathway maintains muscle mass by regulating protein translation through the Akt-mTOR pathway (Fig. 8).

**Figure 8.**
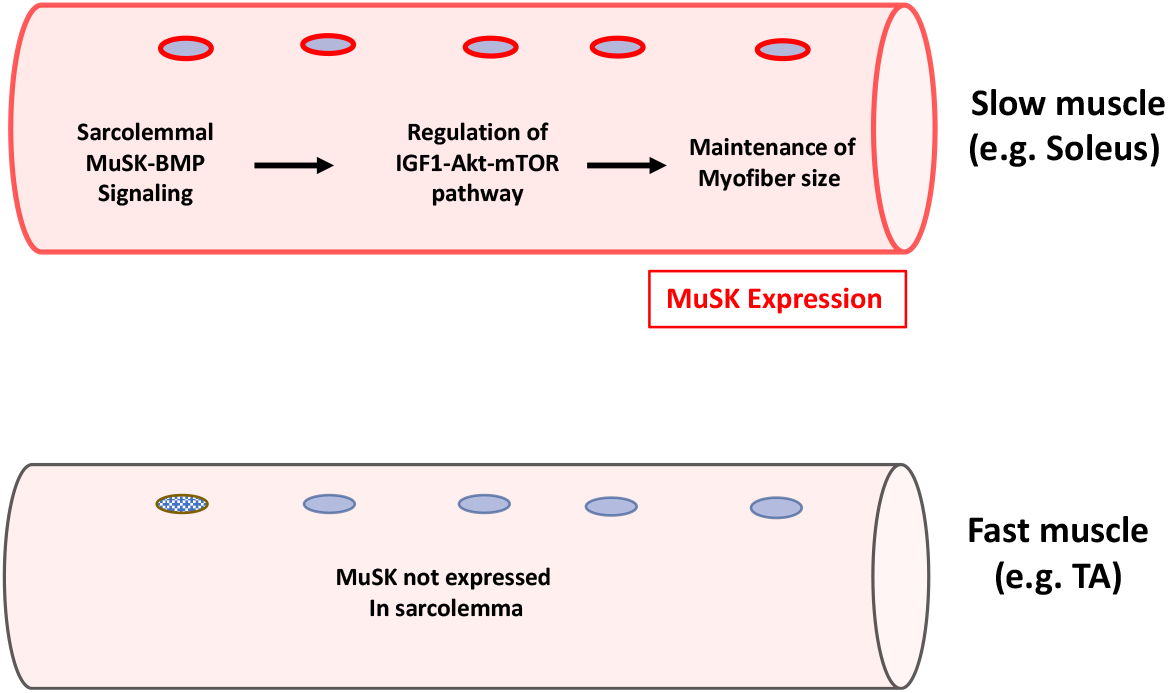
A model of MuSK-BMP regulation of muscle fiber size in slow muscle. MuSK is a BMP co-receptor that is expressed in the sarcolemma (extrajunctionally) in the slow soleus muscle but not in fast muscles such as the TA or EDL. We propose that MuSK-BMP signaling in the sarcolemma selectively regulates myofiber size in slow muscle by controlling the IGF1-Akt-mTOR pathway.

The striking selectivity of the MuSK-BMP pathway in the soleus as compared to the TA is likely to reflect the distinct MuSK expression levels and regulation in this muscle. MuSK transcript levels are ∼4 to 5-fold higher in the WT soleus compared to the fast EDL (Fig. 6). MuSK is localized at NMJs in all muscles^29,32^, including TA and soleus (Fig. 6). However, in WT soleus MuSK is also localized at extrasynaptic domains along the extent of the myofiber (Fig. 6), in agreement with earlier reports^13,16^. Importantly, both MuSK transcript levels and the localization of MuSK at extrasynaptic domains are selectively reduced in ΔIg3-MuSK soleus. This reduction seems likely to be the result of perturbed autoregulation since MuSK itself was identified as a MuSK-BMP dependent transcript in our earlier studies^13^. On the organismal level, our results suggest that MuSK expression in the sarcolemma may be one mechanism conferring muscle-selective regulation of myofiber size in health and disease.

Our results add a novel dimension to our understanding of the role of BMP signaling in regulating muscle size. Previous studies have shown that increasing BMP signaling by overexpression of BMP7 or constitutively active BMPR1a (ALK 3) causes hypertrophy. Notably, the hypertrophy is blocked by the mTOR inhibitor rapamycin, establishing a link between BMP signaling and Akt-mTOR-mediated muscle growth^8^. These results also align with our observation that this pathway is an important output of MuSK-BMP signaling. On the other hand, studies of denervation atrophy have demonstrated a prominent role for ubiquitin ligases and protein degradation in this model of acute loss of muscle mass. We did not observe notable changes in atrogenes in our RNA-seq analysis (Table S4, S5), further supporting the hypothesis that the MuSK-BMP pathway works predominantly via anabolic protein synthesis pathways.

The role of MuSK in maintaining muscle size has potential implications for myasthenia gravis (MG) caused by autoantibodies to MuSK (‘MuSK-MG’)^33^. This form of MG is distinct from the more common anti-AChR MG, can be more severe and does not respond to cholinesterase inhibitors. The pathogenesis of MuSK-MG is mediated at least in part by IgG4 antibodies directed against the MuSK Ig1 domain that disrupt agrin-LRP4 binding and signaling^15,34–37^.

However, some clinical features of MuSK-MG suggest that non-synaptic pathology mediated by the MuSK autoantibodies may also contribute to the disease. MuSK-MG pathology is often more pronounced in restricted muscle groups, including bulbar and respiratory muscles.

Moreover, muscle atrophy is observed in MuSK-MG where it is associated with non-fluctuating weakness, fatty tissue infiltration and myopathic changes in electrophysiology recording. It is therefore plausible that antibodies targeting the MuSK expressed in the sarcolemma could contribute to MuSK MG pathology.

The MuSK-BMP pathway could also be a target for promoting muscle growth and treating conditions associated with muscle atrophy such as sarcopenia, immobilization, and cachexia. Maintenance of muscle mass is a balance between the homeostatic mechanisms regulating protein synthesis and degradation. Although the role for IGF1 as an anabolic pathway is well established, circulating IGF1 levels correlate incompletely with muscle status. Rather, muscle-derived IGF1 is likely to be the dominant mediator of growth^30^. The MuSK-BMP pathway represents an attractive target for developing specific agents to modulate muscle growth. This pathway also offers the prospect of precise manipulation of BMP signaling in muscle. BMPs and their canonical receptors are ubiquitous and manipulating them leads to unwanted side effects; in contrast, MuSK expression is highly enriched in muscle. Moreover, the MuSK ectodomain would be accessible to manipulation by therapeutic antibodies, while antisense oligonucleotides could promote MuSK-BMP signaling without affecting the role of MuSK in synapse formation. Finally, MuSK is expressed in myonuclei in both fast and slow myofibers and its level increases with age in humans and rats^17,38,39^. The MuSK-BMP pathway thus emerges as an attractive target for selectively modulating muscle growth and combatting atrophy.

## MATERIALS AND METHODS

### Animals

To target the MuSK Ig3 domain, we deleted exons 6 and 7 using CRISPR-Cas9 (Exon numbering according to ENSMUST00000081919; Fig. 1) in C57BL/6 mice. Founder mice carrying the MuSK^ΔIg3^ allele were backcrossed to wild type (WT) C57BL/6 background mice, and confirmation of germline transmission of the MuSK^ΔIg3^ to progeny was performed by PCR using isolated genomic DNA. PCR for WT and ΔIg3-MuSK was performed using DreamTaq Green PCR Master Mix (Thermo Scientific) according to manufacturer’s directions using 60°C annealing temperature and 60 second extension for 33 cycles. WT MuSK primers used were forward: TGGGCACTCAATCCAGCAG and reverse: TGGCTAAGCAAGGCAGGAC. ΔIg3-MuSK primers used were forward: TGGGCACTCAATCCAGCAG and reverse: TGGTATCCATCACTTGAACAAG. Offspring were backcrossed to WT C57BL/6 for several generations. Mice of three months of age were used for all experiments. All protocols were conducted under accordance and approval of the Brown University Institutional Animal Care and Use Committee.

### Generation of immortalized ΔIg3-MuSK myogenic lines

WT and ΔIg3-MuSK mice were crossed with the H-2Kb-tsA58 transgenic Immortomouse mouse line (JAX Cat# 032619) obtained from the Jackson Laboratory and immortalized monoclonal myoblast cell lines were established as described previously^40,41^. Myoblasts were isolated from postnatal day 2-3 old pups and cells were subcloned on Matrigel for 1-2 passages, then moved to gelatin coated plates for all future passages^42^. Only clones that were WT and ΔIg3-MuSK homozygous containing the H-2Kb-tsA58 transgene were maintained. Cells were cultured at 33°C in 8% CO2 in Dulbecco’s modified Eagle’s medium supplemented with 20% fetal bovine serum, 2% L-glutamine, 1% penicillin-streptomycin, 1% chicken embryo extract, and 1 U of γ-interferon. For primary cultures, myoblasts were isolated as described previously^42^ and maintained in growth media consisting of Iscove’s modified Dulbecco’s medium (IMDM) containing 20% FBS, 1% CEE and 1% penicillin-streptomycin and differentiated in IMDM supplemented with 2% horse serum and 1% penicillin-streptomycin when myoblasts reached 70% confluency. For BMP4 treatments, cells were serum starved 5 to 6 hr (myoblasts) or were maintained in low-serum differentiation conditions for 24-48 hours (myotubes), then treated with 20-25 ng/ml recombinant human BMP4 protein (R&D Systems, Cat# 314-BP). For acetylcholine receptor clustering experiments, primary myotubes were stimulated with 10 units of agrin (R&D Systems, Cat # 550-AG) for 16 hours and stained with rhodamine-conjugated α-bungarotoxin (Thermo Fisher) at 1:2000. AChR clusters were counted as previously described^43^.

### Histology, immunohistochemistry, and immunocytochemistry

Muscles were flash frozen in freezing isopentane, embedded in Optimum Cutting Temperature media and cryosectioned at 10 μm. For histological analysis sections were stained with Hematoxylin and Eosin. To delineate muscle cell membrane and ECM, sections were stained with Wheat germ agglutinin (WGA) conjugated to Alexa-Fluor 488 in accordance with Treat NMD Standard Operating Procedures (SOP ID MDC1A_M.1.2.002). For muscle fiber sizing experiments, sections of WT and ΔIg3-MuSK TA and soleus muscles were rehydrated and blocked in 20% goat serum then incubated with anti-dystrophin diluted to 1:400 overnight and detected with goat anti-rabbit IgG Alexa Fluor 488 (Invitrogen) secondary antibodies. To identify type I and IIa muscle fibers sizes, a cocktail of anti-dystrophin and mouse IgG1 monoclonal SC-71 (type IIa) or IgG2b BA-D5 (type I) used at 1:100 and detected with fluorescent serotype specific secondaries. Fiber sizes were measured using the open source Quantimus plugin^44^. Muscle sections were stained for MuSK using an IgG4 serum fraction (1:1000) from MuSK myasthenia gravis patients containing MuSK autoantibodies targeting the MuSK Ig1 domain^45^ (a gift from M. Huijbers, Leiden University). Sections were fixed with 1% PFA, washed, and blocked with 2% BSA and 5% goat serum in PBS. Serum was diluted 1:1000 in 1/10 block overnight at 4°C and detected using goat anti-human IgG Alexa Fluor 555 (Invitrogen) diluted 1:1000 and counterstained with AlexaFluor488-conjugated α-bungarotoxin (Thermo Fisher) at 1:1000. WT and ΔIg3-MuSK myoblasts were stained for phospho-Smad1/5 (Ser463/465; CST #9516) according to the manufacturer’s protocol. For MuSK staining in cells, unpermeabilized myoblasts were fixed in 1% paraformaldehyde and immunolabeled with human anti-MuSK monoclonal antibodies targeting the MuSK Ig2 domain (gift from K. O’Connor, Yale University ^46^ and detected with goat anti-human IgG AlexaFlour 555.

Images were acquired on a Nikon Ti2-E inverted microscope equipped with a Photometrics Prime 95B sCMOS camera for fluorescence imaging and a 16-megapixel Nikon DS-Ri2 color camera for imaging histology slides. Confocal z-stack images were obtained using a Zeiss 800 LSM laser scanning microscope equipped with a USRB laser module and GaAsP detectors. When comparing fluorescence levels, all images were acquired on the same session and imaging parameters. Images were processed using ImageJ (NIH).

### Western Blotting

Samples were homogenized in ice-chilled RIPA buffer (Thermo Scientific), cOmplete® EDTA-free protease and phosphatase inhibitor cocktail (Roche) supplemented with 1 mM sodium orthovanadate, 10 mM sodium fluoride, and 1 mM EDTA. Homogenates were centrifuged at 10,000 x g for 10 minutes at 4°C. Total protein concentration of clarified supernatants was determined using the Bradford reagent (Thermo Scientific) and BSA standard (Thermo Scientific). Equal amounts of protein were separated by SDS-PAGE using 4-15% gels (Bio-Rad) and transferred to nitrocellulose membranes using a tank transfer system. Total protein loading for transfer efficiency and normalization was assessed using No-Stain protein labeling reagent (Invitrogen). Membranes were blocked in 5% nonfat dry milk and probed with primary antibodies overnight according to manufacturer’s recommended protocols. All primary antibodies used were rabbit monoclonals obtained from Cell Signaling Tech: phospho-Smad1/5 (Ser463/465) (#9516), total Smad1 (D5957, #6944), phospho-4E-BP1 (Thr37/46) (236B4; #2855), and total 4E-BP1 (53H11, #9644). Primary antibodies were detected with goat anti-rabbit Fc fragment specific HRP-conjugated IgG (Jackson ImmunoResearch, cat. # 111-035-046) used at 1:5000. Membranes were visualized by chemiluminescence and under epi-blue to visualize total protein No-Stain labeling using a ChemiDoc MP imaging system (Bio-Rad). Densitometric analysis was performed using ImageLab software (Bio-Rad).

### Quantitative RT-PCR

Total RNA from snap frozen muscles and cells stored in RNAlater (Invitrogen) was isolated using the RNeasy Fibrous and RNeasy mini kit (Qiagen), respectively, and DNase I treated according to manufacturer’s protocol. cDNA from total RNA was reverse transcribed using SuperScript III cDNA synthesis kit (Invitrogen) and analyzed by quantitative RT-PCR using TaqMan assays (Applied Biosystems) targeting MuSK (Mm00437762_m1), Wnt11 (Mm00437327_g1), Car3 (Mm00775963_g1), and Id1 (Mm001281795_m1). Data analyzed by ΔΔCt method using β-2 microglobulin (Mm00437762_m1) and 18S (Hs99999901_s1) reference genes.

### RNA-seq Analysis

Total RNA from WT and ΔIg3-MuSK mouse soleus and TA was extracted using a RNeasy Fibrous Tissue mini kit (Qiagen) for non-stranded RNA library preparation and sequencing (Genewiz).

Samples were sequenced at a depth of approximately 50 million reads/ sample. Reads were assessed for quality and trimmed of adapter sequences using Trimmomatic then aligned with GSNAP to the Ensembl mouse reference genome (mm10) and read count matrices were generated using htseq-count. WT and ΔIg3-MuSK read count matrices were uploaded to the integrated Differential Expression and Pathway (iDEP) analysis tool for exploratory data analysis, differential gene expression using DEseq2, and gene set enrichment pathway GO analysis (Ge et al., 2018). The raw datasets will be available in GEO immediately following publication.

### Statistical analysis

The average of biological replicates is shown as mean ± SEM. Experiments were replicated two to three times as indicated. Statistical comparisons between groups were performed using Two-way ANOVA and unpaired Student’s t-test when comparing multiple groups or two groups respectively. ANOVA analyses were corrected by post-hoc test as indicated. Significance was determined as P < 0.05 (*** P < 0.001, ** P < 0.01, *P < 0.05).

## Acknowledgements

We thank Beth McKechnie for expert technical assistance and Youngwook Ahn of the Brown Transgenic core for generating the ΔIg3-MuSK mice. We also thank K. O’Connor for generously providing the anti-MuSK monoclonal antibody and M. Huijbers for the MuSK-MG patient IgG4 fraction. We thank the Brown University COBRE Center for Computational Biology P20 GM10903, Mouse Transgenic and Gene Targeting Facility (P30 GM103410), and Genomic Facility (P30 RR031153, P20 RR018728 and S10 RR02763), NSF (EPSCoR grant No 0554548). DJ was supported by 4R25GM083270 and 2T32AG041688, LAF by T32 MH20068 and a Carney Institute Graduate Award. JRF was supported by U01 NS064295, R41 AG073144, R21 NS112743, R21 AG073743, and ALS Finding a Cure.

## Author contributions

DJ: Designed the mouse model; designed, executed, and interpreted experiments (Figs. 1-8); wrote manuscript. LF: generated cell lines; designed, executed, and interpreted experiments (Fig. 1, 2). LM: Designed, executed, and interpreted experiments (Fig. 1, 6). ME: Designed, executed, and interpreted experiments (Fig. 1). JRF: Designed and interpreted experiments, wrote manuscript.

